# Radiation-Driven Prediction of Daily Irrigation Demand under Different Electrical Conductivity Scenarios in Greenhouse Tomato

**DOI:** 10.64898/2026.01.23.701235

**Authors:** Lingang Xiao, Yan Ma, Qian Feng, Xingdong Gao, Huifeng Shi, Xia Liu, Yilei Yin

## Abstract

In soilless greenhouse tomato cultivation, daily transpiration and irrigation demand are largely governed by solar radiation, while irrigation-solution electrical conductivity (EC) used for salinity management may further modulate plant water use. This study developed a low-input, radiation-driven modeling approach to predict daily irrigation demand under contrasting water–salt management scenarios. Two tomato cultivars were grown under four treatments: conventional baselines (CK1, CK2) and regulated scenarios combining irrigation volume with solution EC (low-water high-EC, TK; high-water moderate-EC, TC). Daily irrigation volume (I) and drainage were recorded, and daily cumulative radiation (G) was derived from photosynthetically active radiation (PAR). Within each treatment, we compared a radiation-only baseline model with an EC-adjusted model and evaluated predictive performance using 5-fold blocked time-series cross-validation. Results showed strong positive correlations between G and I across all treatments (p < 0.001). The EC-adjusted models achieved cross-validated root-mean-square errors (RMSE) of 0.815–1.393 L d^−1^ per trough and Nash–Sutcliffe efficiencies (NSE) of 0.407–0.730. Incorporating EC yielded a small but consistent improvement under the TK scenario (ΔRMSE = −0.014 L d^−1^; ΔNSE = +0.019), whereas its effect was negligible or slightly negative under CK1, CK2, and TC, highlighting scenario dependence. Our radiation-driven framework, with an optional EC correction, offers a practical and scalable tool for daily irrigation forecasting and supports integrated water–salt management in soilless greenhouse tomato production.

## 1. Introduction

### 1.1. Greenhouse tomato production and irrigation challenges

Tomato (*Solanum lycopersicum* L.) is a globally significant protected horticultural crop, whose economic viability hinges on yield stability, fruit quality, and resource-use efficiency [1,2]. With the expansion of greenhouse cultivation, management strategies have evolved from a sole focus on yield toward integrated approaches that harmonize productivity, quality enhancement, and water conservation [3,4]. Nevertheless, greenhouse tomato systems remain heavily reliant on irrigation and fertigation, rendering water management a critical—and often limiting—factor in the context of escalating water scarcity and rising production costs [5–7].

In practice, irrigation scheduling in greenhouses frequently depends on grower experience or fixed timetables, which seldom account for rapid daily fluctuations in crop water demand. Suboptimal irrigation not only curtails water-use efficiency but also perturbs root-zone salinity and nutrient dynamics [8,9], thereby influencing photosynthetic performance, yield formation, and key fruit quality attributes such as soluble solids and nutritional composition [10–12]. Hence, developing irrigation strategies that dynamically respond to environmental variability while safeguarding yield and quality remains a pivotal challenge in modern greenhouse tomato production.

### 1.2. Radiation-driven irrigation demand in greenhouse systems

Among environmental drivers, solar radiation exerts a dominant influence on canopy transpiration and carbon assimilation in greenhouse tomatoes [13,14]. Daily variations in photosynthetically active radiation (PAR) directly modulate stomatal conductance and transpiration rates, establishing radiation as a primary determinant of short-term irrigation demand [15–17]. Unlike temperature and humidity, which can be partially regulated in controlled environments, incident radiation is largely governed by external weather conditions, leading to pronounced day-to-day variability in crop water requirements [18,19].

Empirical relationships between daily cumulative radiation and irrigation water consumption have been consistently demonstrated in greenhouse tomatoes, furnishing a practical foundation for radiation-based estimation and scheduling approaches [13,20,21]. Compared with mechanistic evapotranspiration models or data-intensive machine-learning techniques, empirical radiation-driven models offer distinct advantages in simplicity, robustness, and operational feasibility—attributes especially valuable for daily irrigation scheduling in commercial settings [22–25]. However, most existing radiation-based studies have been conducted under uniform fertigation regimes, seldom explicitly addressing potential interactions with root-zone salinity management [26].

### 1.3. Electrical conductivity and water–salt interactions

In soilless and substrate-based greenhouse tomato systems, the electrical conductivity (EC) of the nutrient solution is a key operational variable that shapes the root-zone water–salt environment [27–29]. Adjustments in EC alter osmotic potential and ion concentrations around roots, thereby influencing water uptake, transpiration, and assimilate partitioning [30,31]. Numerous studies indicate that moderate elevation of EC, or controlled deficit irrigation, can enhance fruit quality traits such as soluble solids and sugar-acid ratio, though excessive salinity may suppress yield and overall plant water consumption [32–34].

Critically, the effects of EC on crop water use are not independent of irrigation supply level. Under differing water-availability backgrounds, identical EC conditions may exert contrasting influences on transpiration and irrigation demand, owing to shifts in plant water status and root-zone hydraulic properties [35,36]. Recent work emphasizes the importance of integrated water–salt regulation rather than treating EC as a static background parameter [37–40]. Despite this, most irrigation-scheduling frameworks continue to neglect EC-dependent adjustments in daily irrigation demand, particularly in soilless cultivation systems.

### 1.4. Research gap and objectives

Although radiation-driven approaches are widely employed to estimate greenhouse irrigation demand, prevailing frameworks rarely incorporate EC-related adjustments under contrasting water–salt management scenarios. This omission constrains their applicability for integrated irrigation strategies that simultaneously aim for water conservation and fruit quality improvement.

Therefore, this study aimed to develop a simple, practical radiation-based method for daily irrigation demand estimation under varying electrical conductivity scenarios in substrate-grown greenhouse tomato. Specific objectives were to:(i) quantify the relationships between daily cumulative radiation and irrigation demand under different EC management backgrounds;(ii) elucidate how EC modifies radiation-driven water-demand responses; and (iii) evaluate the implications of EC-adjusted irrigation scheduling for yield stability and fruit quality.

By emphasizing operational feasibility and employing treatment-wise blocked time-series cross-validation, this work provides a management-oriented irrigation framework that complements more complex modeling approaches reported in the literature.

## 2. Materials and Methods

### 2.1. Experimental Site and Environmental Conditions

The experiment was conducted from November 2023 to May 2024 in Glass Greenhouse A2 at the Facility Agriculture Research Base of the Academy of Agricultural Planning and Engineering, Ministry of Agriculture and Rural Affairs, located in Yongqing County, Langfang City, Hebei Province, China. The greenhouse covered approximately 5000 m^2^ and was equipped with automated environmental-control systems (internal/external shading, fan-pad cooling, heating, supplemental lighting, and ventilation). The daytime temperatures were controlled between 22°C and 26°C, and night-time temperatures ranged from 18°C to 22°C. Relative humidity was maintained at 60%-70%, and environmental data (such as radiation and temperature) were continuously recorded using an on-site automatic weather station.The complete daily environmental monitoring and irrigation records are provided in S1 Dataset.

### 2.2. Plant Materials and Cultivation Method

Test varieties were ‘Jingdan No. 8’ (cherry tomato) and ‘Jingcai No. 8’ (strawberry tomato). Soilless substrate cultivation used a 3:1 (v/v) coconut coir:perlite mix. Each substrate trough measured 1.0 m × 0.15 m × 0.12 m (length × width × height), with four plants per trough (plant spacing 30 cm, row spacing 55 cm) (Figure 1). Drip irrigation was applied using municipal tap water, with nutrient solution adjusted for pH and EC before irrigation. Prior to planting, coconut coir was fully hydrated and pretreated with standard nutrient solution to minimize initial water-salt variability.

**Figure 1.**
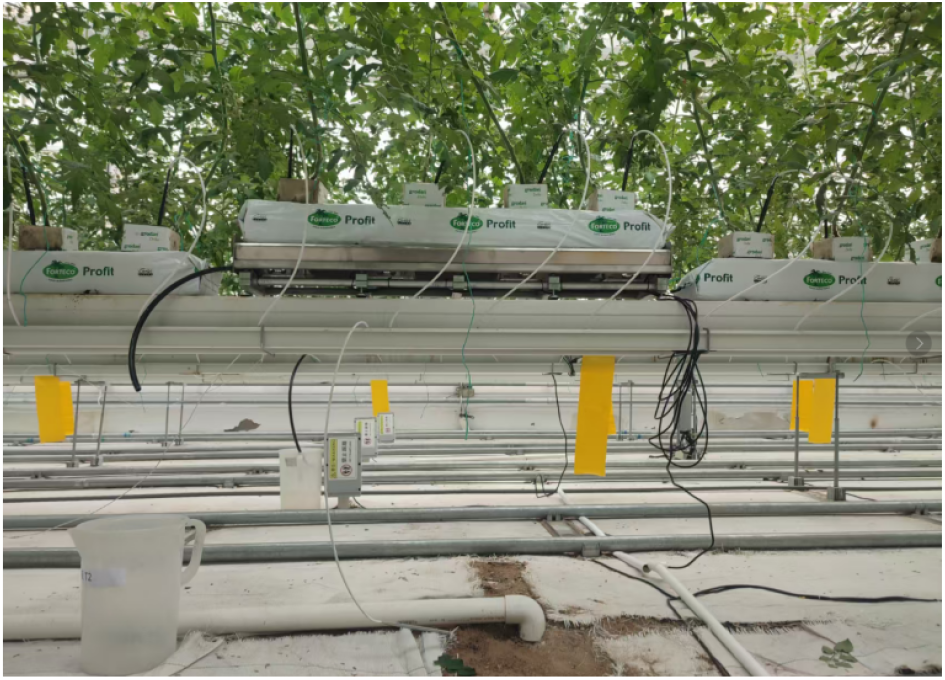
Schematic diagram of the substrate cultivation trough and drip irrigation setup for greenhouse tomato experiments.

### 2.3. Experimental Design and Treatments

A randomized block design was used with four water-fertilizer treatments: conventional water-salt baseline for each variety (CK1 and CK2) and corresponding regulated scenarios (low-water high-EC: TK; high-water moderate-EC: TC). Each treatment had three replicates (12 plots total).

For model development, daily measurements were averaged across the three replicate troughs within each treatment, yielding one daily observation per treatment per day (approximately 180 days per treatment).

Considering varietal differences in water demand and nutrient management, a “variety–water-salt scenario pairing” framework was adopted. ‘Jingdan No. 8’ received conventional baseline (CK1) and low-water high-EC (TK) to analyze supply level and EC interactions under the same variety. ‘Jingcai No. 8’ received conventional baseline (CK2) and high-water moderate-EC (TC) to examine EC adjustments under enhanced supply. CK1 and CK2 served as practical baselines for their respective varieties. Although initial design intended pH adjustment for CK2, actual irrigation pH remained similar across CK1 and CK2 (Table 1); thus, CK2 was positioned as the baseline for ‘Jingcai No. 8’ without treating pH as an independent factor.

**Table 1.**
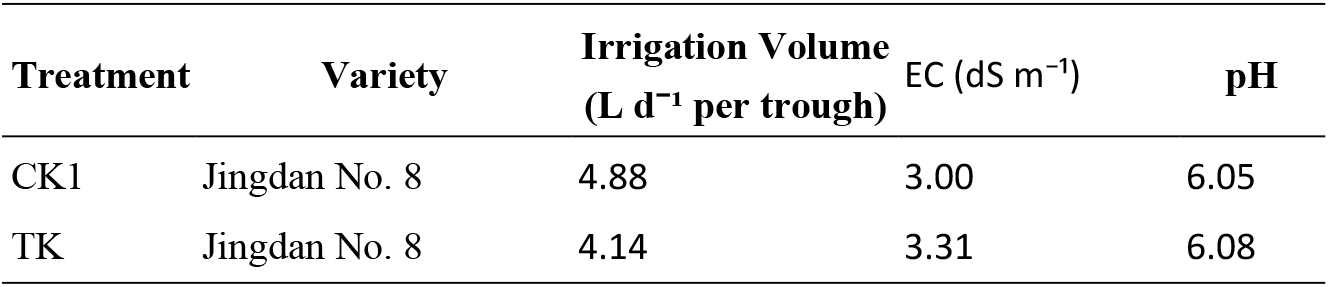
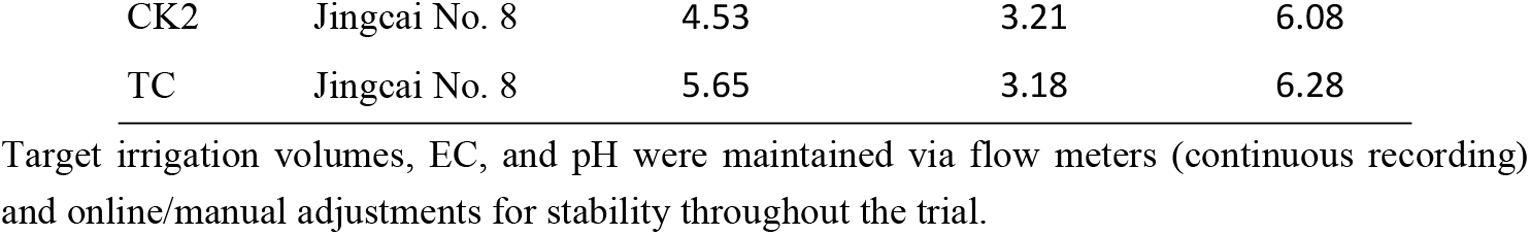
Experimental treatment setup.

| Treatment | Variety | Irrigation Volume<br>(L d<sup>-1</sup> per trough) | EC (dS m<sup>-1</sup>) | pH |
| --- | --- | --- | --- | --- |
| CK1 | Jingdan No. 8 | 4.88 | 3.00 | 6.05 |
| TK | Jingdan No. 8 | 4.14 | 3.31 | 6.08 |
| CK2 | Jingcai No. 8 | 4.53 | 3.21 | 6.08 |
| TC | Jingcai No. 8 | 5.65 | 3.18 | 6.28 |

Target irrigation volumes, EC, and pH were maintained via flow meters (continuous recording) and online/manual adjustments for stability throughout the trial.

### 2.4. Measurements and Data Collection

Irrigation volume was recorded in real time by flow meters; drainage was collected and summarized daily. Nutrient solution EC and pH were measured using conductivity (INESA DDS-307A) and pH meters (PHS-3C), calibrated periodically.

Light conditions were monitored with quantum sensors (LI-190R, LI-COR, USA) placed at greenhouse top, recording PAR at 1-min intervals. Daily cumulative radiation (G, MJ m^−2^ d^−1^) was integrated from PAR. Specifically, PAR (μmol m^−2^ s^−1^) was integrated to daily light integral (DLI, mol m^−2^ d^−1^) and converted to energy units using a conversion factor of 0.218 MJ per mol photons. At the daily aggregation scale used for modeling, no missing values were present; therefore, no interpolation or exclusion was required. As a prespecified quality-control rule, if PAR gaps had occurred, gaps ≤10 min would have been linearly interpolated, whereas days with >5% missing PAR would have been excluded from modeling.

Photosynthetic parameters (net photosynthetic rate Pn, transpiration rate Tr, stomatal conductance Gs, intercellular CO_2_ concentration Ci) were measured using a portable system (LI-6400XT) on three representative plants per treatment (three measurements per plant, averaged). Photosynthesis measurements are available in S4 Dataset.

Yield was recorded at the plot level (one cultivation trough per replicate) at each harvest (fruit number and total fresh weight). Yield was expressed as g plot^−1^ per harvest (and kg plot^−1^ for seasonal cumulative yield). Per-harvest yield and fruit number data are available in S2 Dataset.

### 2.5. Data Processing and Statistical Analysis

Data were organized in Microsoft Excel and analyzed using SPSS (version 25.0). Treatment effects were assessed via one-way ANOVA at α = 0.05, with Duncan’s multiple range test applied for post-hoc comparisons. Pearson correlation coefficients were computed to evaluate linear associations, with significance levels denoted as *p < 0.05, **p < 0.01, and ***p < 0.001.

To quantify the combined influence of radiation and irrigation-solution EC on daily irrigation volume, an EC-adjusted linear regression model was fitted separately for each treatment:

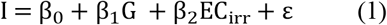

A radiation-only baseline model was also fitted for comparison:

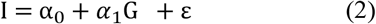

where I is daily irrigation volume (L d^−1^ per trough), G is daily cumulative radiation (MJ m^−2^ d^−1^) integrated from PAR, EC_irr_ is the electrical conductivity of the irrigation solution (dS m^−1^), β_i_ and *α*_i_ are regression coefficients, and ε is the random error term.

Prior to final model fitting, a 3σ residual rule was applied within each treatment to exclude outliers: an initial model was fitted, residuals were calculated, and observations with |residual| > 3σ were removed. The model was then refitted on the filtered dataset.

To evaluate out-of-sample predictive performance while accounting for temporal autocorrelation, a blocked 5-fold time-series cross-validation was implemented. Daily observations were ordered chronologically and partitioned into five contiguous time blocks. In each fold, one block served as the validation set, while the remaining four blocks constituted the training set. The 3σ outlier removal was applied solely to the training data within each fold. Predictions from all folds were pooled, and performance metrics—root-mean-square error (RMSE), mean absolute error (MAE), and Nash–Sutcliffe efficiency (NSE)—were computed from the pooled out-of-sample predictions.

## 3. Results

### 3.1. Correlations among Key Irrigation–Drainage Variables

Daily-scale relationships among irrigation, drainage, and radiation variables were examined using Pearson correlations, including irrigation volume (I), irrigation-solution electrical conductivity (EC_irr_) and pH, drainage volume (D), drainage EC (EC_dra_) and pH, and daily cumulative radiation (G). The correlation structure is illustrated in Figure 2 **(underlying data in S1 Dataset)**.

**Figure 2.**
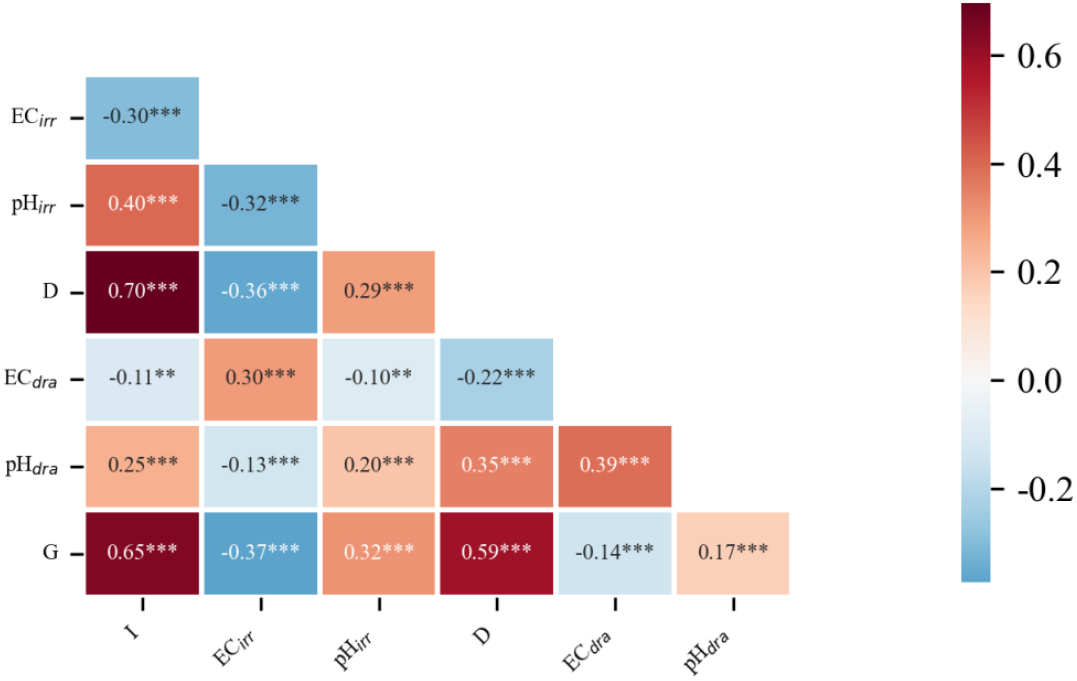
Correlation matrix of key irrigation, drainage, and radiation variables at the daily scale. Color intensity indicates correlation strength and direction. Asterisks denote significance (* p < 0.05, ** p < 0.01, *** p < 0.001). I, irrigation volume (L d^−1^ per trough); D, drainage volume (L d^−1^ per trough); G, daily cumulative radiation (MJ m^−2^ d^−1^); EC_irr_, electrical conductivity of irrigation solution (dS m^−1^); pH_irr_, pH of irrigation solution; EC_dra_, electrical conductivity of drainage solution (dS m^−1^); pH_dra_, pH of drainage solution.

I was strongly and positively correlated with D (r = 0.70, p < 0.001), consistent with the coupled dynamics of substrate water input and outflow. G was also positively correlated with I (r = 0.65, p < 0.001), supporting the dominant role of radiation-driven transpiration in determining day-to-day irrigation demand.

Among solution properties, EC_irr_ was moderately negatively correlated with irrigation pH (r = −0.32, p < 0.001). In addition, G was negatively correlated with EC_irr_ (r = −0.37, p < 0.001), suggesting that EC tended to be reduced on high-radiation days, potentially as an operational adjustment to mitigate salt accumulation risk.

### 3.2. Daily Irrigation Demand Prediction Models

Daily-scale empirical regression models were developed separately for CK1, CK2, TK, and TC, using daily accumulated radiation (G) and irrigation water electrical conductivity (EC_irr_) as predictors and daily irrigation amount (I) as the response variable.

Drainage electrical conductivity (EC_dra_) was not statistically significant during candidate-variable screening (p > 0.05) and was therefore excluded from the final models.

CK1:

I = 2384.62 + 2065.40G ‐ 421.35EC_irr_

R^2^ = 0.79, n = 179; G showed a highly significant positive effect (p < 0.001), while EC_irr_ showed a significant negative effect (p = 0.025).

CK2:

I = 605.17 + 1687.12G + 259.51EC_irr_

R^2^ = 0.71, n = 180; G showed a highly significant positive effect (p < 0.001). The effect of EC_irr_ was positive but marginal (p = 0.052).

TK:

I = 1252.60 + 995.08G + 360.98EC_irr_

R^2^ = 0.52, n = 180; G showed a highly significant positive effect (p < 0.001), and EC_irr_ showed a significant positive effect (p = 0.032).

TC:

I = 3511.60 + 1555.54G ‐ 191.25EC_irr_

R^2^ = 0.61, n = 182; G showed a highly significant positive effect (p < 0.001). The effect of EC_irr_ was negative but not significant (p = 0.279).

Overall, the four models consistently indicate that G is the dominant driver of day-to-day irrigation variation, whereas the direction and significance of EC_irr_ depend on the water management scenario.

### 3.3. Model Performance Evaluation

Model performance was evaluated using the coefficient of determination (R^2^), root mean square error (RMSE), mean absolute error (MAE), and Nash–Sutcliffe efficiency (NSE) (Table 2). RMSE and MAE quantify daily scheduling errors in physical units (L d^−1^ per trough), whereas NSE summarizes how well the model reproduces day-to-day variability relative to the observed mean.

**Table 2.** Performance indicators of radiation-driven daily irrigation models under different treatments.

| Treatment | n | R<sup>2</sup> | RMSE ($\text{L d}^{-1}$ per trough) | MAE ($\text{L d}^{-1}$ per trough) | NSE |
| --- | --- | --- | --- | --- | --- |
| CK1 | 179 | 0.79 | 1.01 | 0.79 | 0.79 |
| CK2 | 180 | 0.71 | 0.93 | 0.72 | 0.71 |
| TK | 180 | 0.52 | 0.75 | 0.57 | 0.52 |
| TC | 182 | 0.61 | 1.13 | 0.85 | 0.61 |

Across treatments, the fitted radiation-driven models achieved RMSE values of 0.75–1.13 L d^−1^ per trough and MAE values of 0.57–0.85 L d^−1^ per trough (Table 2), indicating that daily irrigation demand was typically estimated within approximately 0.6–0.9 L d^−1^. For the in-sample fitted models, NSE values numerically matched R^2^ because both metrics were computed as 1 − SSE/SST on the same dataset with an intercept term; therefore, RMSE and MAE are particularly informative for operational scheduling accuracy.

Performance differed among water–salt scenarios. CK1 exhibited the strongest fit (R^2^/NSE = 0.79), suggesting a stable radiation–irrigation response under conventional management for ‘Jingdan No. 8’. CK2 and TC showed intermediate explanatory power (R^2^/NSE = 0.71 and 0.61, respectively), whereas TK had the lowest explained variance (R^2^/NSE = 0.52), implying larger residual variability under the low-water high-EC regime. Notably, TK also showed the smallest absolute errors (RMSE = 0.75 L d^−1^; MAE = 0.57 L d^−1^), consistent with the lower irrigation supply level limiting the magnitude of day-to-day fluctuations (Table 1).

In relative terms, RMSE corresponded to approximately 18–21% of the mean daily irrigation volume across treatments, supporting the practical interpretability of the error magnitudes for daily scheduling. Because in-sample metrics may overstate predictive performance when temporal dependence is present, an out-of-sample assessment based on blocked time-series validation is reported in the following section.

### 3.4 Baseline Comparison and Blocked Validation

Within each treatment, a radiation-only baseline model was compared with an EC-adjusted model using blocked 5-fold time-series cross-validation. Daily observations were ordered chronologically and split into five contiguous time blocks; in each fold, one block was held out for validation and the remaining blocks were used for training. Validation predictions were pooled across folds, and RMSE, MAE, and NSE were computed from the pooled out-of-sample predictions (Table 3).

**Table 3.** Blocked 5-fold time-series cross-validation performance of the radiation-only baseline model and the EC-adjusted model (pooled across folds). **Note:** RMSE and MAE are expressed in L d^−1^ per trough; NSE is dimensionless. Δ indicates

| Treat<br>ment | n | RMSE<br>(baseline) | MAE<br>(baseline) | NSE<br>(baseline) | RMSE<br>(EC-adjusted) | MAE<br>(EC-adjusted) | NSE<br>(EC-adjusted) | ΔRMSE<br>(EC – baseline) | ΔMAE (EC – baseline) | ΔNSE<br>(EC – baseline) |
| --- | --- | --- | --- | --- | --- | --- | --- | --- | --- | --- |
| CK1 | 179 | 1.110 | 0.905 | 0.744 | 1.141 | 0.926 | 0.730 | 0.031 | 0.021 | -0.014 |
| CK2 | 180 | 1.020 | 0.800 | 0.655 | 1.032 | 0.811 | 0.647 | 0.012 | 0.011 | -0.008 |
| TK | 180 | 0.829 | 0.629 | 0.422 | 0.815 | 0.607 | 0.441 | -0.014 | -0.022 | 0.019 |
| TC | 182 | 1.313 | 0.955 | 0.473 | 1.393 | 0.991 | 0.407 | 0.080 | 0.036 | -0.066 |

EC-adjusted minus baseline; negative ΔRMSE/ $\Delta$ MAE and positive ΔNSE indicate improvement.

The cross-validated results indicate that the incremental value of adding EC was scenario dependent. Under the low-water high-EC scenario (TK), incorporating EC reduced RMSE from 0.829 to 0.815 L d^−1^ per trough and increased NSE from 0.422 to 0.441 (ΔRMSE = −0.014; ΔNSE = +0.019), reflecting a small but consistent improvement. Although the magnitude of improvement was modest, the direction of change (ΔRMSE < 0 and ΔNSE > 0) suggests that EC carries incremental predictive information only under specific water–salt backgrounds. In contrast, EC adjustment slightly degraded performance under CK1 and CK2 (ΔNSE = −0.014 and −0.008, respectively) and produced a larger decline under TC (ΔRMSE = +0.080; ΔNSE = −0.066). These comparisons support treating EC as a scenario-specific adjustment factor—most useful under low-water high-EC management—rather than a universal predictor that improves accuracy across all treatments.

Compared with in-sample fitting, NSE decreased under cross-validation, as expected when predicting unseen time blocks, with baseline NSE ranging from 0.422 to 0.744 across treatments. Pooled observed–predicted relationships corroborate the numerical comparison: predictions broadly followed the 1:1 reference across treatments, with a modest reduction in scatter for TK under the EC-adjusted model and substantial overlap between the two models for CK1, CK2, and TC (Figure 3). Collectively, the blocked validation confirms that cumulative radiation captures the dominant day-to-day signal, while the additional contribution of EC is scenario dependent.

**Figure 3.**
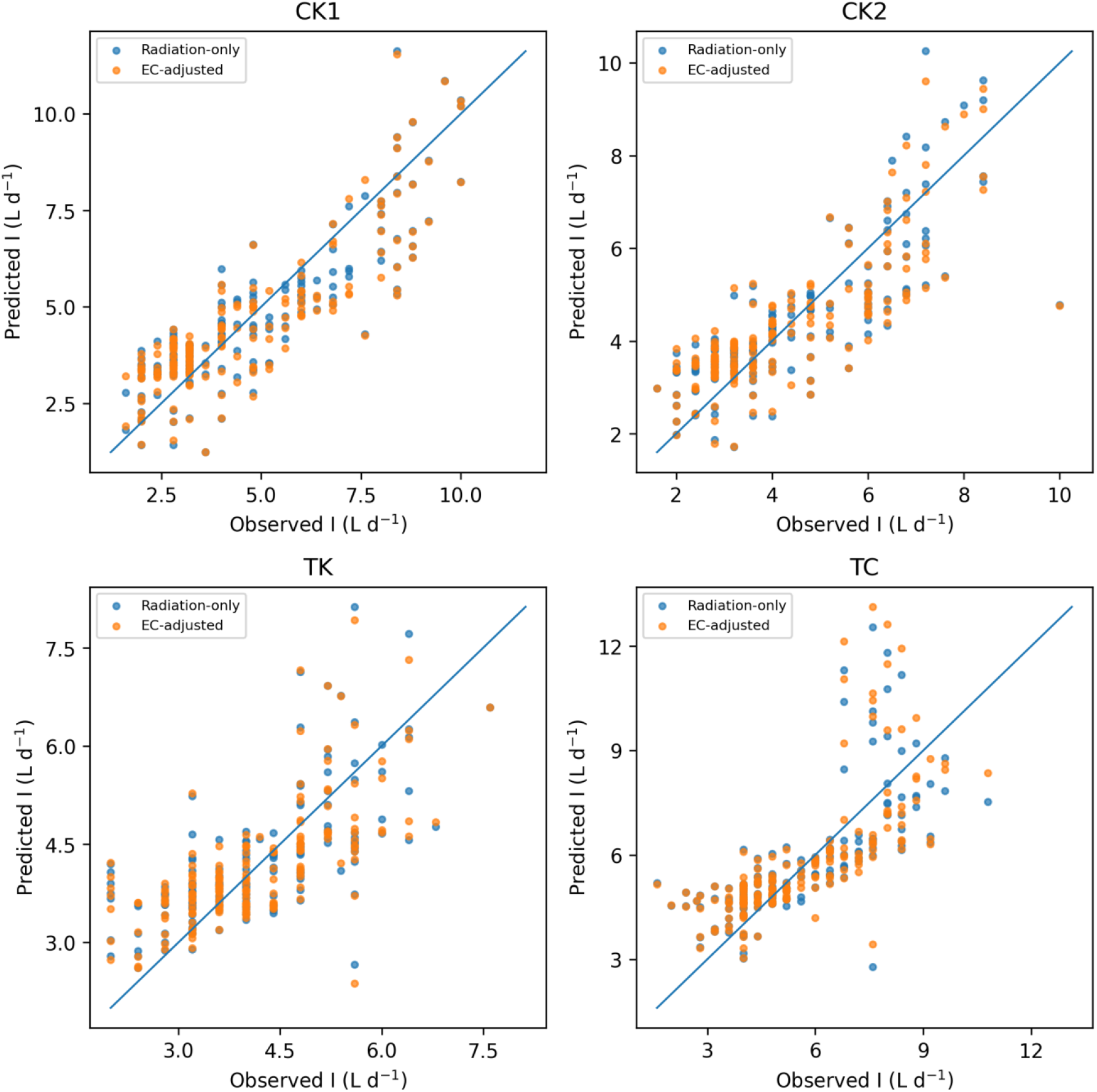
Observed versus predicted daily irrigation volume (I, L d^−1^ per trough) for the blocked 5-fold time-series cross-validation (pooled predictions) under each treatment. Predictions from both the radiation-only baseline model and the EC-adjusted model are shown. The solid line indicates the 1:1 reference line.

### 3.5. Radiation-Driven Irrigation and Drainage Responses

Across treatments, G maintained a highly significant positive association with I (p < 0.001), and the strength of the relationship varied by scenario. The strongest G–I correlations occurred under CK1 (r = 0.73) and TC (r = 0.72) (Figure 4), reinforcing the consistency of radiation-driven water demand under contrasting management backgrounds.

**Figure 4.**
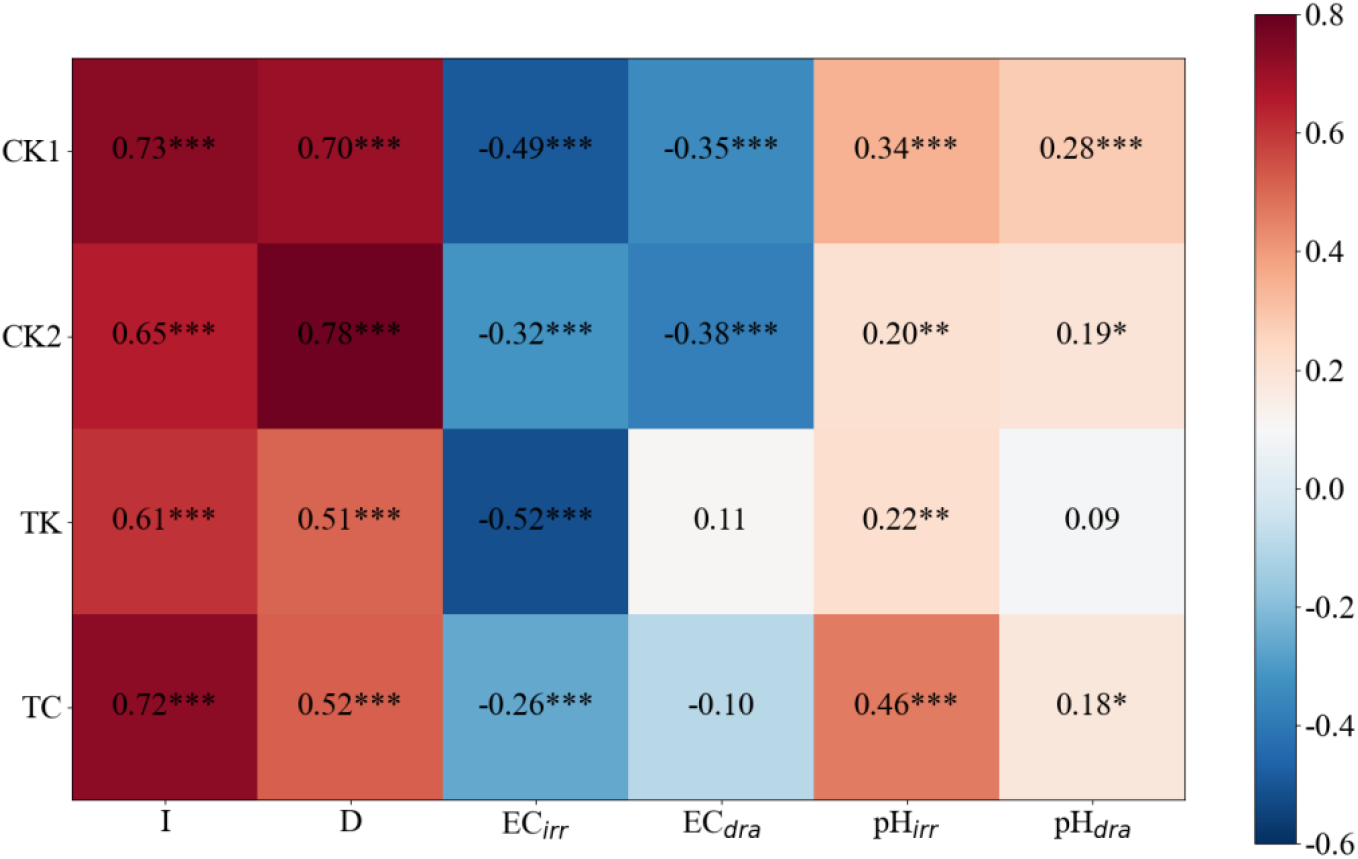
Correlation analysis between daily cumulative solar radiation (G, MJ m^−2^ d^−1^) and irrigation/drainage variables (I and D, L d^−1^ per trough) under different treatments.

Relationships between G and drainage volume were more treatment sensitive. The G–D correlation was highest under CK2 (r = 0.78), followed by CK1, while weaker responses were observed under TK and TC (r ≈ 0.51–0.52). This pattern suggests that drainage reflects not only radiation-driven irrigation input but also scenario-dependent root uptake and substrate buffering effects.

Correlations between G and solution properties were generally weaker. Associations with EC_irr_ and pH were mostly small (|r| < 0.35), with significant negative G–EC_irr_ correlations only under specific treatments. Relationships involving irrigation and drainage pH were weaker still, indicating that daily variations in EC and pH were primarily governed by nutrient-solution preparation and substrate buffering rather than direct radiation forcing.

### 3.6. Yield Dynamics

Fresh fruit yield per harvest (g plot^−1^) exhibited pronounced seasonal variation (Figure 5; S2 Dataset). Low yields in early winter were followed by increasing harvest amounts toward spring, with peaks in April–May as radiation and temperature improved.

**Figure 5.**
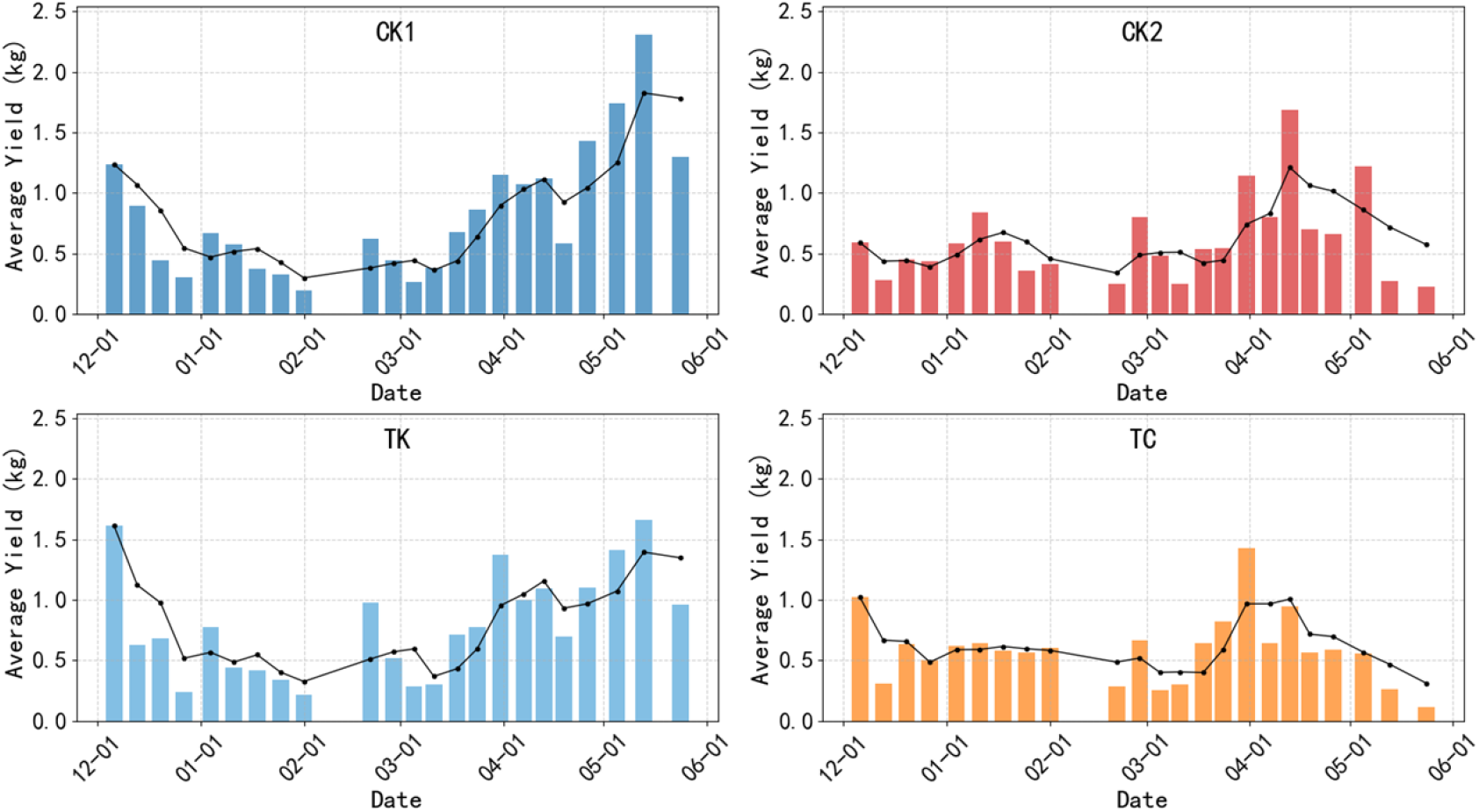
Temporal variation in fresh fruit yield (g plot^−1^ per harvest) under different treatments.

For ‘Jingdan No. 8’ (CK1 and TK), cumulative yield increased steadily throughout the season. For ‘Jingcai No. 8’ (CK2 and TC), yield remained lower early in the season and increased later, indicating more stage-dependent responses under the corresponding water–salt regimes.

Late-season dynamics differed between the two cultivar–scenario groups. ‘Jingcai No. 8’ displayed greater fluctuations: CK2 peaked in mid-April (1.68 kg d^−1^), whereas TC peaked in late March (1.43 kg d^−1^), followed by sharp declines in late May (CK2: 0.23 kg d^−1^; TC: 0.12 kg d^−1^). In contrast, ‘Jingdan No. 8’ maintained comparatively better late-season performance under both CK1 and TK.

### 3.7. Fruit Nutritional Quality

Fruit quality was assessed using ascorbic acid/vitamin C (AsA/VC, mg 100 g^−1^ FW) and sugar–acid ratio (SAR = TSS/TA) under contrasting seasonal radiation conditions (S3 Dataset), representing low-light winter (January) and high-light spring (April). Treatment distributions in the AsA/VC–SAR space shifted markedly between these periods (Figure 6), with lower values in January and higher values in April, consistent with improved nutritional quality and flavor balance under increased radiation.

**Figure 6.**
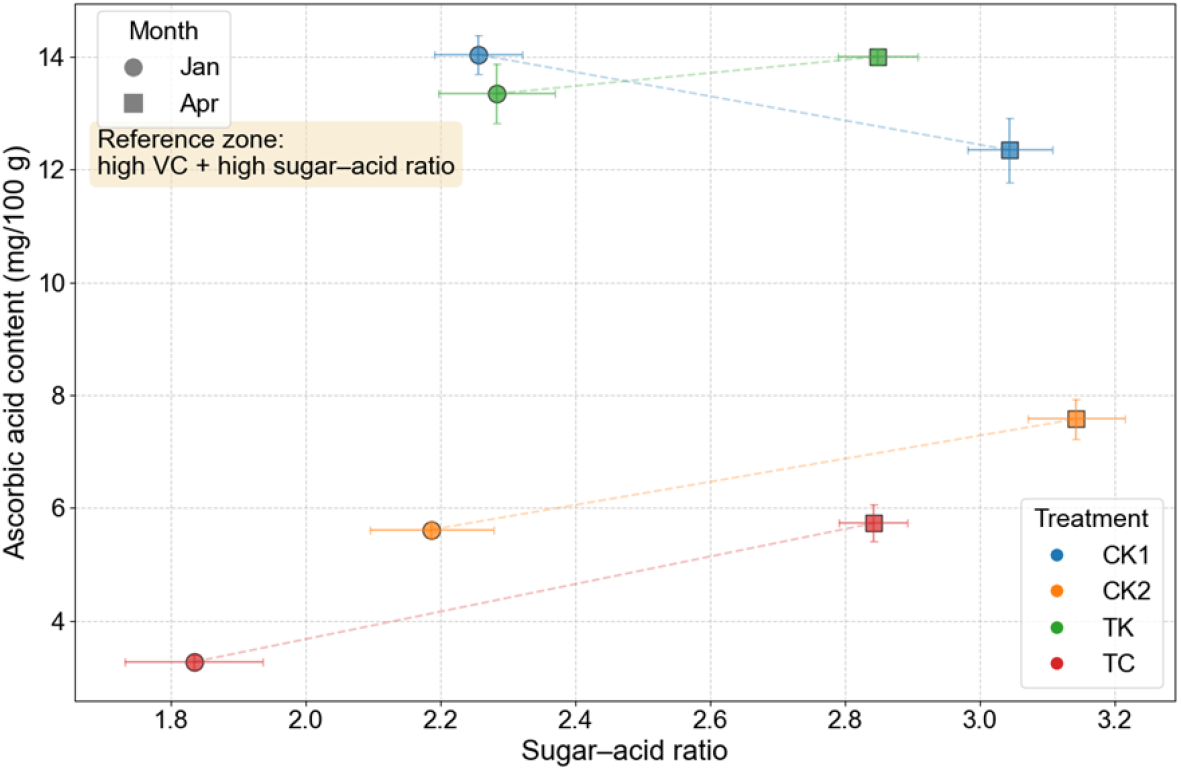
Fruit quality expressed as ascorbic acid (AsA/VC, mg 100 g^−1^ FW) versus sugar–acid ratio (SAR = TSS/TA, dimensionless) under different treatments. TSS is in °Brix and titratable acidity (TA) is in g 100 g^−1^ FW (citric acid equivalents). The boundary of “high VC + high SAR” is empirical for reference only.

Within ‘Jingdan No. 8’, CK1 maintained relatively high AsA/VC but lower SAR than TK. For ‘Jingcai No. 8’, AsA/VC was generally lower across treatments, and TC remained consistently low. Treatment separation became more apparent under higher radiation, suggesting that light availability modulated the expression of water–salt management effects on fruit composition.

### 3.8. Seasonal Photosynthetic Responses

Seasonal changes in photosynthetic traits are summarized in Figure 7 (S4 Dataset). Net photosynthetic rate (Pn) increased from January to May across treatments. In January, TK showed the highest mean Pn (13.18 μmol CO_2_ m^−2^ s^−1^), while TC was lowest (7.99 μmol CO_2_ m^−2^ s^−1^). In May, mean Pn values converged across treatments (13.51–14.14 μmol CO_2_ m^−2^ s^−1^), with CK1 slightly higher than the others.

**Figure 7.**
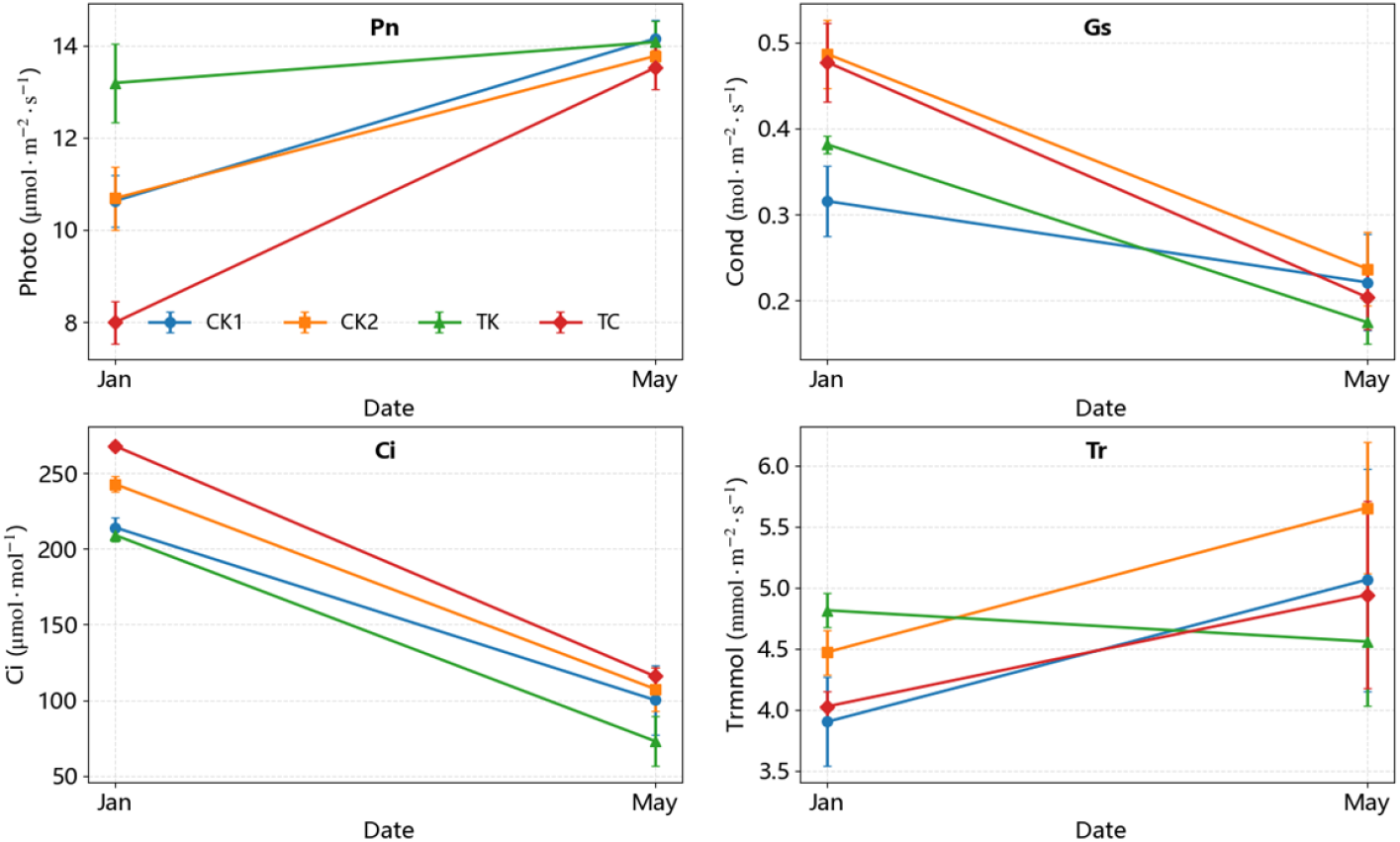
Temporal variation of photosynthetic parameters (Pn, Gs, Ci, and Tr; units shown on axes) in tomato under different treatments.

Stomatal conductance (Gs) generally decreased from January to May, accompanied by a decline in intercellular CO_2_ concentration (Ci). Transpiration rate (Tr) increased seasonally in CK1, CK2, and TC, whereas TK maintained relatively higher Pn with comparatively lower Tr, indicating scenario-specific decoupling between carbon assimilation and water loss.

## 4. Discussion

Limitation (study design): treatments were cultivar-bound (CK1-TK for ‘Jingdan No. 8’; CK2-TC for ‘Jingcai No. 8’), so treatment contrasts are interpreted within each cultivar-specific case study and we avoid cross-cultivar causal claims.

### 4.1. Radiation as the Primary Driver of Daily Irrigation Demand

Daily cumulative solar radiation (G) showed consistently strong and highly significant positive correlations with irrigation volume across all treatments, confirming radiation-driven transpiration demand as the dominant determinant of short-term water requirements in soilless greenhouse tomatoes [13,33]. This finding aligns with previous studies demonstrating that, under protected cultivation, daily water uptake is more tightly coupled to incident radiation than to air temperature or humidity alone, particularly when canopy development and root-zone moisture are maintained within suitable ranges [5,11–14]. Radiation governs stomatal opening, leaf energy balance, and transpiration flux, thereby directly translating light variability into irrigation demand at the daily scale [30–32].

Compared with approaches based on stage-based coefficients or evapotranspiration estimates requiring multiple environmental inputs, the radiation-driven relationship observed here provides a simplified yet robust basis for irrigation scheduling [10,15]. Within each cultivar-specific treatment set (CK1–TK for ‘Jingdan No. 8’ and CK2–TC for ‘Jingcai No. 8’), the G–I relationship remained stable, suggesting that radiation is the primary driver under the management backgrounds examined. Accordingly, G can be used as the core predictor for daily irrigation decision-making, while EC and irrigation volume are incorporated as scenario-specific adjustments rather than as cross-cultivar general rules.

### 4.2. Scenario-Dependent Effects of Electrical Conductivity on Irrigation Demand

Although radiation dominated daily irrigation variability, the role of irrigation solution electrical conductivity (EC) exhibited clear scenario dependence. Under conventional water supply (CK1), EC showed a significant negative association with irrigation demand, suggesting that higher salinity increased osmotic resistance in the root zone, thereby suppressing plant water uptake [41]. In contrast, under the low-water high-EC scenario (TK), EC exerted a positive effect on irrigation demand, indicating that moderate salinity stress may have enhanced transpiration or induced compensatory water uptake responses under limited water supply [27–29].

These contrasting responses highlight that EC effects should not be interpreted independently of irrigation background. Under sufficient water availability, elevated EC primarily increases osmotic stress and reduces effective water absorption [21,22]. Under restricted irrigation, however, higher EC may alter root hydraulic conductivity, stomatal regulation, or assimilate partitioning in ways that partially offset water limitation [20, 23]. Similar interaction effects between salinity and irrigation regime have been reported in greenhouse tomatoes, where moderate salinity improved certain physiological or quality traits without proportionally reducing water uptake [42].

This scenario dependence also helps interpret why EC improved blocked cross-validation only under TK. One likely reason is that EC_irr_ was operationally adjusted in response to radiation (significant negative G–EC_irr_ correlations occurred only under specific treatments), which can introduce collinearity and temporal non-stationarity that weakens out-of-sample generalization in CK1/CK2/TC. Under TK, where irrigation supply is restricted and salinity management is more tightly coupled to plant water status, EC_irr_ may better reflect osmotic constraints affecting water uptake and thus provides incremental predictive signal beyond radiation.

Collinearity between G and EC_irr_ was mild in all treatments (pairwise r = −0.26 to −0.52; VIF = 1.07–1.36), suggesting that the scenario-dependent gain/loss is more likely driven by how EC was operationally adjusted over time than by severe multicollinearity.

Consistent with this interpretation, under the high-water moderate-EC scenario (TC), EC effects on irrigation demand were weak and non-significant, indicating that abundant water supply can buffer salinity-related impacts on short-term water demand [8,9]. Together, these results emphasize that EC should be treated as a scenario-adjustment parameter rather than a universal modifier in radiation-driven irrigation models.

### 4.3. Performance and Practical Applicability of Radiation-Driven Empirical Models

The empirical regression models based on daily radiation and EC achieved moderate to high explanatory power (R^2^ = 0.52–0.79) across treatments, with RMSE values of 0.75–1.13 L d^−1^ and NSE values closely aligned with R^2^ [16–18,35,36]. These performance metrics indicate that the models effectively captured major day-to-day fluctuations in irrigation demand, while prediction errors were primarily associated with occasional extreme deviations rather than systematic bias [13,17].

It is important to emphasize that the proposed models are intended as applied scheduling tools rather than mechanistic simulations of plant or root-zone processes [10,34]. By relying on easily measurable inputs and treatment-specific coefficients, the models balance accuracy with operational feasibility, which is a key consideration for greenhouse management [11,15]. Similar empirical or semi-empirical approaches have been widely adopted in protected horticulture to support real-time decision-making, particularly where full physiological modeling is impractical [12,16,18].

The blocked cross-validation results further confirm that daily cumulative radiation is the dominant predictor of day-to-day irrigation demand, as the radiation-only baseline already provided robust accuracy across treatments. The additional EC term did not uniformly improve cross-validated errors, but its contribution was clearly scenario-dependent, consistent with the direction and significance of EC effects observed in the fitted models. Therefore, EC is better interpreted as a scenario-adjustment factor for water–salt management rather than a universal accuracy booster.

### 4.4. Yield and Quality Responses under Integrated Water–Salt Management

Seasonal yield dynamics reflected the combined influences of radiation availability, temperature, and water–salt management [2,3,33]. Low yields during winter coincided with limited radiation, while yield peaks in spring corresponded to improved light conditions [1,13]. Under the low-water high-EC treatment (TK), yield accumulation was initially comparable to or slightly higher than the conventional treatment but declined later in the season, indicating stage-specific sensitivity to water–salt stress [27,28].

Quality responses showed clearer differentiation among treatments. Higher ascorbic acid content and sugar–acid ratios under TK, particularly during high-radiation periods, suggest that moderate water and salinity stress promoted assimilate concentration and quality enhancement [43]. This trade-off between yield stability and quality improvement is consistent with previous findings that controlled stress can enhance fruit quality attributes in greenhouse tomatoes while maintaining acceptable yield levels [22,23,29,44].

Notably, quality differences among treatments became more pronounced under higher radiation, indicating that light availability modulates the expression of water– salt management effects on fruit composition [30–32]. These results underscore the importance of explicitly integrating radiation conditions when evaluating irrigation and salinity strategies aimed at balancing yield and quality.

### 4.5. Variety-Specific Responses, Limitations, and Management Implications

Varietal responses differed under comparable management frameworks. ‘Jingdan No. 8’ exhibited greater yield stability and quality potential under both conventional and low-water high-EC conditions, whereas ‘Jingcai No. 8’ showed weaker late-season performance under high-water moderate-EC management [2,3,45]. These differences likely reflect varietal variation in root architecture, salinity tolerance, and photosynthetic regulation, reinforcing the need for variety-specific optimization rather than universal irrigation or EC targets [8,9].

Several limitations should be acknowledged. Treatments were variety-bound, which restricts direct cross-variety comparisons of EC effects. In addition, pH variation among treatments was limited, preventing robust evaluation of pH–water– salt interactions [7,39]. Future studies should incorporate factorial designs with clearer gradients of EC and pH within the same variety, as well as mixed-effects modeling approaches to better address heterogeneity [16,17].

From a management perspective, the results suggest that radiation-driven irrigation prediction can serve as the core component of daily scheduling, with EC used to adjust strategies according to water availability and varietal characteristics [10,11,15]. Such an approach provides a practical pathway toward integrated water– salt management in soilless greenhouse tomato production.

## 5. Conclusions

1. Daily irrigation demand in soilless greenhouse tomato was primarily driven by cumulative solar radiation (G), showing a strong positive association with irrigation amount (I) and drainage (D). A radiation-only linear baseline therefore provides a parsimonious predictor for day-to-day irrigation forecasting.
2. In 5-fold blocked time-series cross-validation, the radiation-driven models achieved RMSE of 0.815–1.393 L d^−1^ per trough and NSE of 0.407–0.730 across treatments. Adding irrigation-solution electrical conductivity (EC_irr_) yielded a small improvement under the low-water high-EC scenario (TK), but changes were negligible or negative under CK1, CK2, and the high-water moderate-EC scenario (TC). Thus, EC should be treated as a scenario-specific adjustment rather than a universal driver.
3. Differences in yield, fruit quality, and photosynthetic traits were consistent with the contrasting water–salt management backgrounds, supporting that irrigation forecasting should be interpreted together with production targets. However, inference remains limited to the cultivar–management combinations tested (CK1–TK in ‘Jingdan No. 8’; CK2–TC in ‘Jingcai No. 8’).
4. For practice, we recommend a two-step scheduling workflow: estimate daily irrigation from G using the radiation-only baseline, then optionally apply an EC-based correction where validated (e.g., low-water high-EC conditions), and distribute the daily total into irrigation events according to greenhouse control settings. This provides a low-input pathway toward integrated water–salt management for soilless greenhouse tomato production.

## Author Contributions

Conceptualization, Y.Y. and Y.M.; Methodology, L.X., Y.M. and Y.Y.; Formal analysis, L.X. and Q.F.; Investigation, X.G., H.S. and X.L.; Data curation, L.X. and X.L.; Writing—original draft, L.X.; Writing—review and editing, Y.Y., Y.M. and Q.F.; Visualization, L.X.; Supervision, Y.Y.; Project administration, Y.M. and Y.Y.; Funding acquisition, Y.Y.

## Data Availability Statement

All relevant data underlying the findings of this study are available within the manuscript and its Supporting Information files. Specifically:

S1 Dataset: Daily environmental parameters (radiation, temperature), irrigation volumes, and drainage records used for modeling.

S2 Dataset: Detailed yield and fruit number data sorted by harvest date.

S3 Dataset: Fruit nutritional quality measurements (Vitamin C, soluble sugar, titratable acidity).

S4 Dataset: Photosynthetic characteristics (Pn, Gs, Ci, Tr) measured at different growth stages.

S1 Code: Python script used for analyzing fruit quality and generating Figure 6. There are no restrictions on data access.

## Ethics statement

Not applicable.

